# The Computational Neuroanatomy of Predictive Dynamics of Pain Perception

**DOI:** 10.1101/2022.04.13.488260

**Authors:** Ryota Ishikawa, Jun Izawa

**Affiliations:** Ph.D. Program in Humanics, University of Tsukuba, Ibaraki 305-8573, Japan; Faculty of Engineering, Information, and Systems, University of Tsukuba, Ibaraki 305-8573, Japan

**Author notes:** Corresponding author: Jun Izawa, Faculty of Engineering, Information, and Systems, University of Tsukuba, 1-1-1 Tennodai, Tsukuba, Ibaraki 305-8573, Japan.

## Abstract

Pain perception is an active process that regulates nociceptive inputs by descending opioidergic signals, in which the brain encodes pain-related predictive and corrective terms, after having made Bayesian-like inferences about noxious amplitudes. Offset analgesia (OA), a large reduction of tonic pain after a small nociceptive termination, is typical empirical evidence of on-line pain modulation through prediction and its correction. However, the basic computational structure underlying OA is not understood. Here, we adopted a constructive approach, formulated the inference of noxious amplitudes with a Kalman filter model, i.e., a recursive Bayesian computation, and then deduced the computational structure for OA, in which an interaction between two latent state variables was implemented. Simulation results suggested that the unidirectional interaction of the two states with two dissociable roles (an integral over time and a derivative of stimulus changes) is crucial for OA. Our results, combined with previous anatomical studies, suggest a computational basis of neural connectivity for pain. The ACC and aINS interact to compute a descending prediction to the brainstem, i.e. PAG, while ascending inputs are filtered in the thalamus and delivered to the cortices as prediction errors. Thus, we suggest dissociable, computational roles of the ACC and aINS in pain processing.

**Author Summary:** Understanding the computational theory of pain perception is crucial for clarifying why some painful syndromes become chronic. Here, we propose a computational neuroanatomical model of endogenous pain modulation and we simulate a model for offset analgesia. We first demonstrate through model comparisons that the brain encodes at least two distinct states to estimate ongoing nociception: a derivative of input changes and its integral. We suggest that its neural substrate comprises hierarchical circuits composed of cortices, the thalamus, and brainstem. Second, we show that the computational basis of disrupted pain modulation in patients is pseudo-neglect of actual sensory inputs, with bias toward the internal prediction. Our results are the first to provide a neurocomputational mechanism of pain perception dynamics and a factor that determines its functionality.

## Introduction

Pain processing is an essential cognitive function for organismal survival. After an injury, persisting tonic pain is an important cue to monitor the condition of the body and to choose appropriate actions, e.g., resting, favoring the injured structure, or escaping. Modern theories have proposed that multiple regions and the network they form (Fig. 1) represent such pain processing [1–3], since there is no unitary region for pain in the brain, i.e., a “pain cortex”[4–6]. However, because of its complexity, the means by which the neural processing of these regions is integrated still remains puzzling.

**Fig.1.**
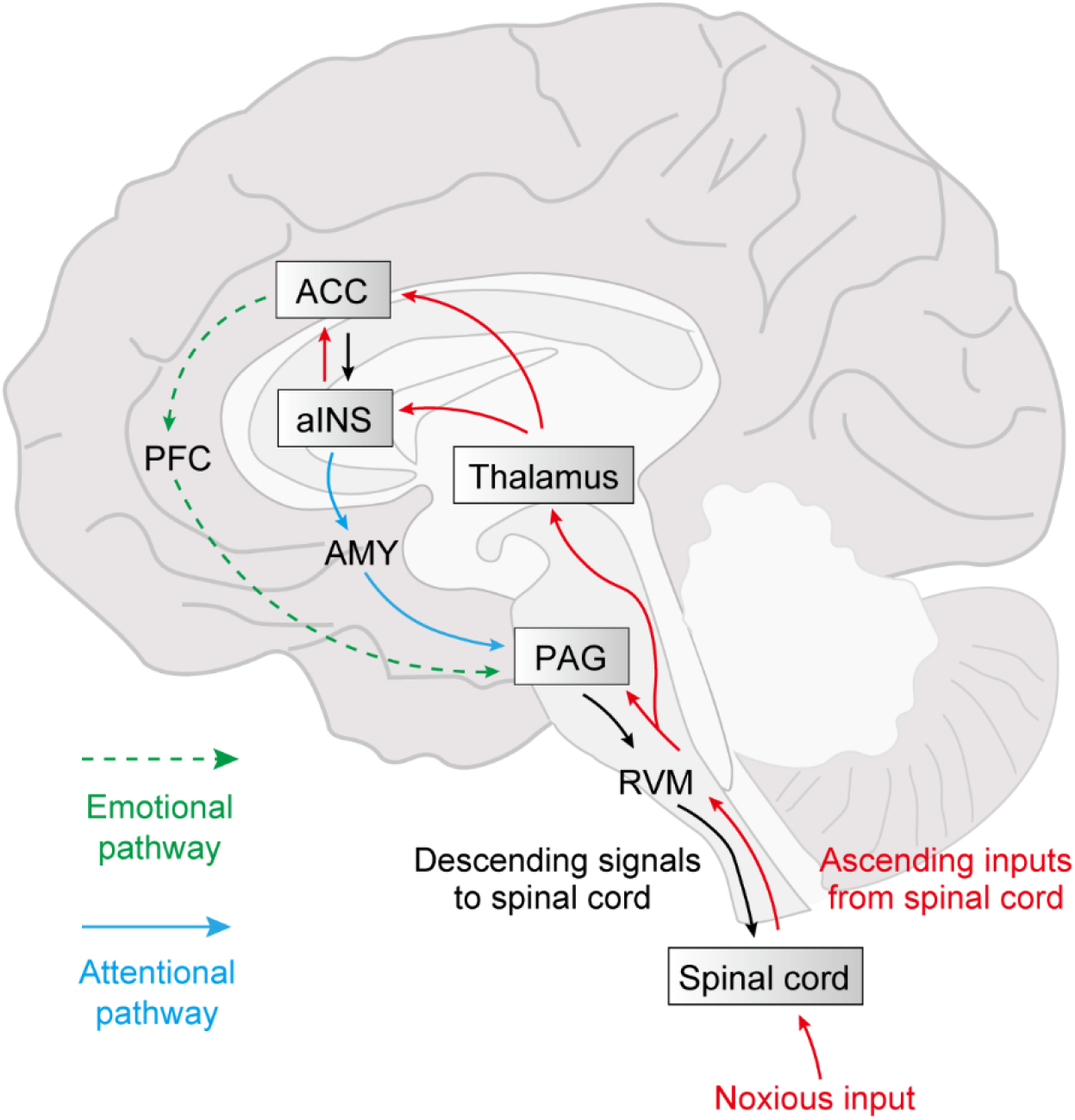
A consensus on neuroanatomy and a pain processing network. The representative theory of the neural mechanism of pain postulates ascending and descending pathways [3,35–37]. The ascending pathway (red arrows) conveys nociceptive inputs from the spinal cord to the cortex via the thalamic nucleus. The descending pathway projects from cortical regions to the brainstem, e.g., the Periaqueductal Gray (PAG) or the Rostral Ventromedial Medulla (RVM), finally arriving at the spinal cord to modulate further afferent inputs (black arrows). It has been thought that in this process there should be two modulatory circuits, i.e., one that consists of the anterior cingulate cortex (ACC) and the prefrontal cortex (PFC), which provide emotional modulation (green dashed arrow), and another that consists of the anterior insula cortex (aINS) and the amygdala (AMY), which supply attentional modulation (cyan solid arrow).

Theoretically, Bayesian computation is fundamental in sensory perception, such as vision and touch [7–11]. Pain perception is also thought to involve Bayesian computation. For instance, various studies have suggested that placebo and nocebo effects for phasic pain, e.g., a pulse of noxious heat, can be explained by Bayesian inference of pain, in which the prediction of nociceptive inputs, i.e., priors, is integrated with the stimulus input, i.e., likelihood, leading to pain perception, i.e., posterior [12–16]. Meanwhile, the current painful experience itself should provide effective cues to expect incoming tonic pain. For example, an increase in pain intensity often signals still more pain to come [17].

Offset analgesia (OA) is well-known empirical evidence of this effect, defined as a disproportionately large reduction of perceived pain intensity immediately after a small, i.e., 1°C decrease in the presented temperature [18–20]. Hyperalgesia, which is induced by the opposite temperature pattern of OA is called onset hyperalgesia (OH), and is thought to have a common neural basis, although there is less empirical and neuroimaging evidence [21–23]. Since these temporal dynamics characterize neural processing tonic pain, illustrating the computational model based on Bayesian inference should be useful to ascertain which neural structures are engaged and how they contribute.

Clinically, OA has been used as an index of endogenous pain modulation, and the deficit of this phenomenon, i.e., less or no analgesic effect after reduction of a noxious heat stimulus, has been reported in patients with neuropathic pain [24,25] and other chronic pain syndromes [26–30]. These dysfunctions can be caused by descending modulatory regions, e.g., the anterior cingulate cortex (ACC) and brainstem, which showed weaker BOLD signals in patients than in healthy controls [31]. This seemed to produce a slower pain perception [28], but few investigations have identified the neuropathological mechanism underlying attenuation of OA in chronic pain patients [32].

To figure out the computational structure underlying OA and OH, we have built a computational model of tonic pain perception, based on a recursive Bayesian computational process, i.e., a Kalman filter [33]. The purpose here is to test various structures of hidden causes of pain, i.e., latent states and noxious intensity, i.e., observable value, in Kalman filter models that are necessary to replicate characteristics of OA and OH reported in human studies (Fig. 2). This approach to computational structure may also reveal corresponding neural structures with dissociable cortical functions [34]. In particular, the total number of hidden variables and how they interact in the identified computational model, provide a basis to understand how many neural areas are involved and how they are connected in the brain. For example, it is widely known that emotion and attention are processed in distinct descending circuits to modulate pain [3,35–37] (Fig. 1). This can be modeled by the two latent states that interact equivalently, or else there is no interaction between them. Meanwhile, the predictive coding framework [38–40] has proposed a hierarchical structure between neural populations to encode the world, i.e., top-down predictive pathways and bottom-up corrective pathways.

**Fig.2.**
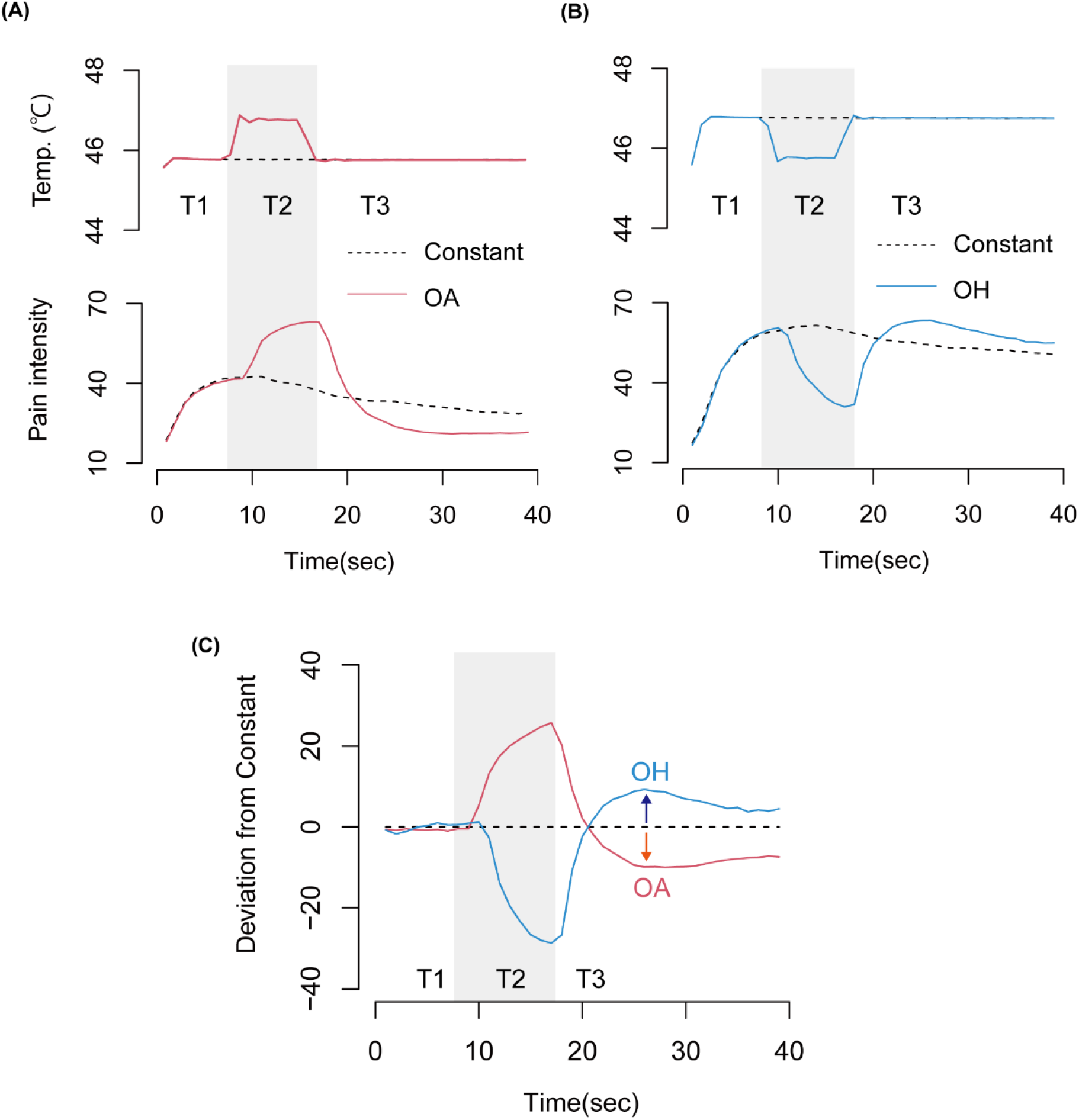
Offset analgesia and onset hyperalgesia. **(A)** The pain intensity rating for the temperature of the offset analgesia (OA) condition (solid red line) showed an increase in T2, but was largely reduced in T3 compared to the constant condition (dashed black line). **(B)** The pain intensity rating for the temperature of the onset hyperalgesia (OH) condition (solid cyan line) decreased in T2, but largely increased in T3 compared to the constant condition (dashed black line). **(C)** Removing the effect of habituation, which resulted in a gradual rating decrease over time in all conditions, deviations of the pain intensity rating of OA and OH from that of the constant condition revealed more clearly the OA and OH phenomena. These temperature patterns and pain intensity ratings were reproduced in accordance with previous literature[22].

Thus, we hypothesized that the brain represents one or more latent state variables that represent an environmental source of pain and that this computational structure is crucial for OA and OH. (1) In order to explain OA and OH, are two latent states necessary or not? (2) If so, how do they generate observable output? (3) How do they interact? (4) How does this theory explain the neural basis of chronic pain? By identifying these structures in a framework of predictive coding with the normative model using Kalman filter theory, we provide insights on the unknown anatomical structure involved in tonic pain processing as a computational constraint on neuroanatomy.

## Results

### General framework

Typical temporal profiles of OA and OH are shown in Figs. 2A-B. During T3 phase, on-line rating of perceived pain intensity was undershot in the OA condition and overshot in the OH condition, compared to the constant baseline condition (Fig. 2C). Here, we aimed to replicate such effects with our computational models. Our model, based on a Kalman filter, comprises two processes: state prediction and state estimation. In the state prediction phase, the latent state in the next time step is predicted by the previous estimate. In the state estimation phase, such a prediction is refined by prediction error, i.e., the difference between measured input and its estimate after filtering, depending on measurement precision.

We test our computational models and examine which structure may explain the properties of human pain perception with the representative experiment paradigm that uses the continuous heat stimuli on the skin, including a slight decrease and increase above the pain threshold (Fig. 3A). The thermal stimulus consisted of three phases: the initial painful stimulus (T1, during 5 sec), 1-2°C increase/decrease to the second stimulus (T2, during 5 sec), and return to the T1 stimulus (T3, during 10 sec). In T3 phase, perceived pain intensity is expected to reduce/increase disproportionately to the presented temperature. Compared to the baseline condition (T1:45-T2:45-T3:45°C, BL), we tested two temperature patterns of (1) OA1: 45-46-45°C and (2) OA2: 45-47-45°C. The OA2 condition should be useful for testing whether the extent of the analgesic effect is proportional to the step size [23], i.e., the step size of +2°C results in a larger analgesic effect than that of +1°C. Furthermore, to account for a bidirectional modulation, i.e., OH, in a unified model, we also decided to test the OH1 condition, (3) OH1: 45-44-45°C, where the perceived pain intensity is expected to increase disproportionately in T3 phase. All of these four stimulus patterns start from 44.5 °C (1sec) and terminate with 44°C (1sec), which is just above the pain threshold, since OA and OH have been discussed as phenomena for painful stimuli, not painless ones.

**Fig.3.**
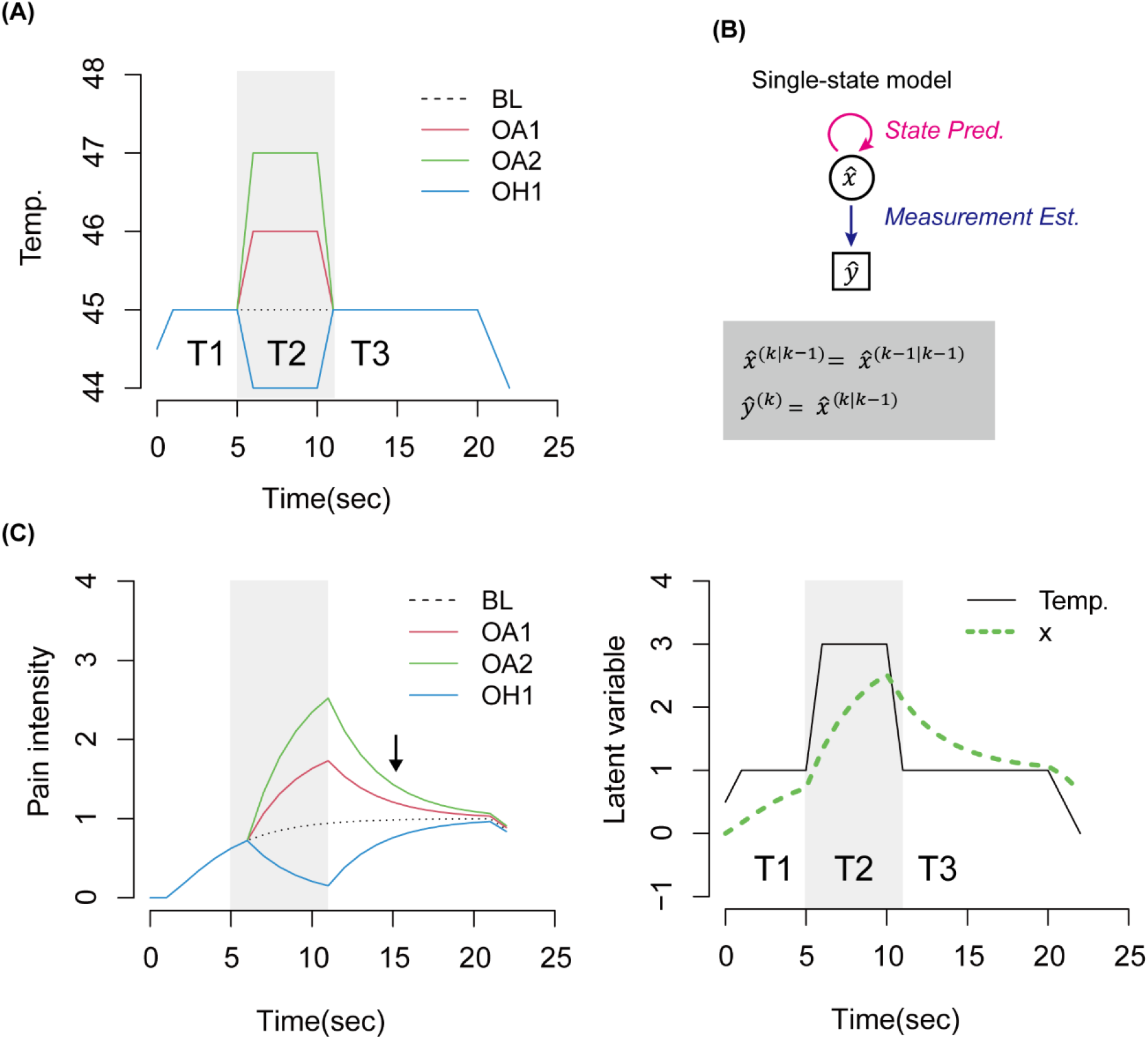
Thermal stimulus and single-state model. **(A)** The thermal stimulus used in the simulations. **(B)** Diagram of single-state model structure. The circle indicates the latent variable, and the square indicates the measurement estimate. The magenta arrow indicates the prediction of the next state. The blue arrow indicates the measurement estimate from the predicted latent variable. The state equation and measurement equation are described within the gray rectangle. **(C)** The left is the simulation result of pain intensity. Simulations under the conditions of four temperature patterns are shown by dotted black, solid red, green, and cyan lines. The black arrow indicates that the single-state model did not induce any effects of OA or OH. The right is the simulation result of a latent state variable under the OA2 condition. The dotted green line indicates the dynamics of the latent variable, and the solid black line indicates the temperature pattern of OA2.

### The single state did not explain OA or OH

We first tested whether the single-state model (Fig. 3B) could replicate OA and OH. The main goals of this simulation were to determine: (1) whether OA1/OA2 conditions produce an undershoot, i.e., analgesia, in T3 phase and (2) whether the OH1 condition produces an overshoot, i.e., hyperalgesia in T3 phase. Briefly, this model failed to produce these phenomena, since there were no undershoots/overshoots in T3 phase. (Fig. 3C, left). For example, in the OA2 condition, the latent state variable of the pain prediction 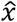 increased in T2 phase, but in T3 phase diminished to the same value as in T1 phase, without any deviation from the presented temperature (Fig, 3C, right). This indicated that although the Kalman filter model with the single state successfully estimated the presented temperature accurately, it did not replicate either OA or OH. These results suggest that more than one state variable in the Kalman filtering model is necessary to reproduce OA and OH phenomena. Results of further model testing with two-state models are shown in the following sections.

### Model comparison within model family 1: the parallel contribution of the state to measurement estimation

In model family 1, where two state variables 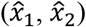 additively generate a prediction of pain intensity, we tested three model structures. The graphical models and corresponding state-space representations of the tested models are summarized in Fig. 4A. In the “No interaction” model (Nint), 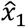 and 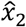 do not interact, i.e., each of them is predicted independently in the next step from its own previous estimate. In the “Unidirectional interaction” model (Uint), 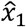 in the next step is updated from both previous estimates of 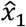 and 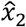, whereas 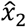 in the next step is updated only on its own. In the “Bidirectional interaction” model (Bint), both variables in the next step are predicted from their own and the other’s previous estimates.

**Fig.4.**
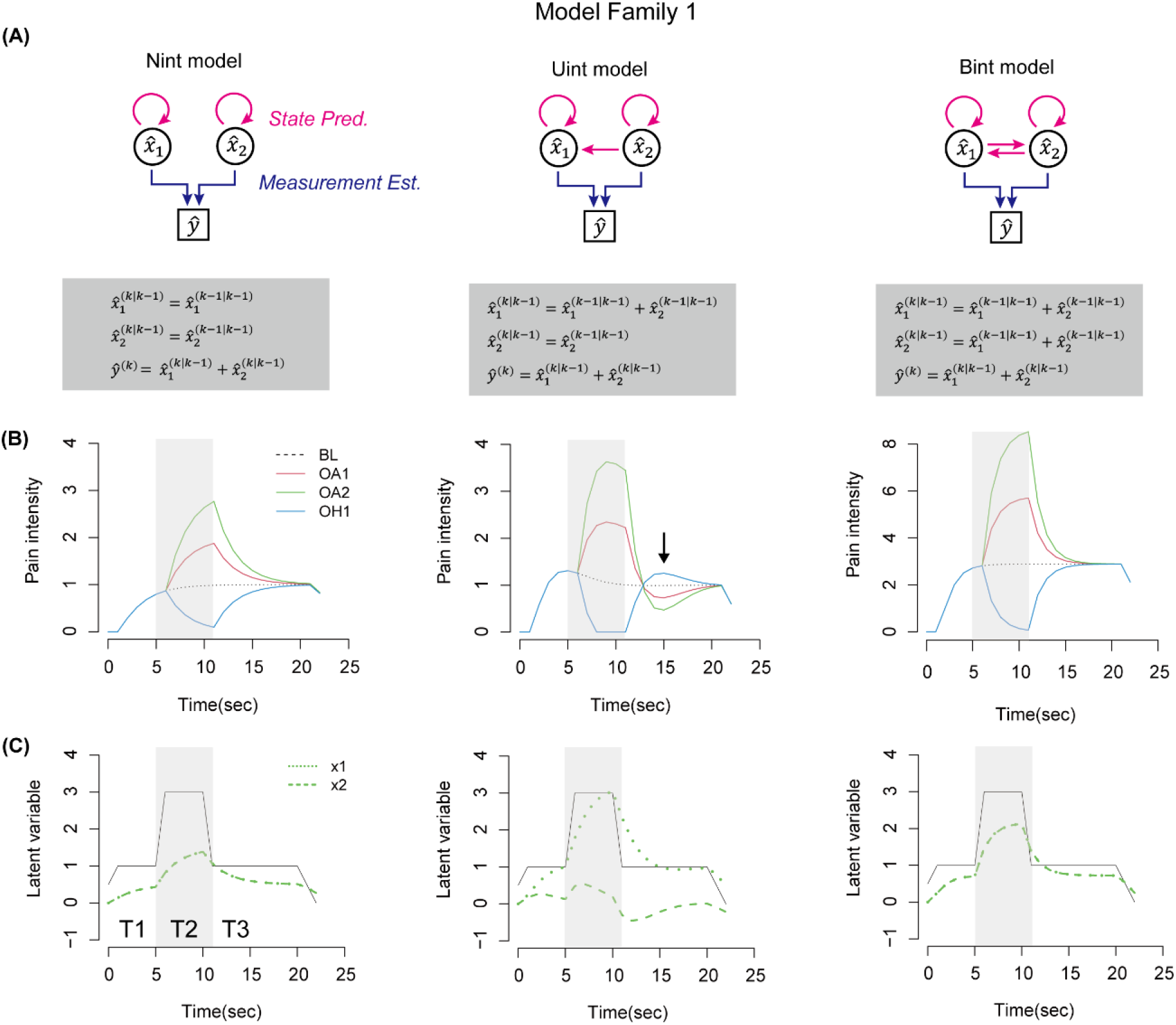
Model family 1. (**A**) Diagrams of each model structure. Circles indicate latent variables, and squares indicate the measurement estimate. Magenta arrows indicate the prediction of the next state. Blue arrows indicate the measurement estimate from latent variables. The state and measurement equations are described in the gray rectangles. (**B**) Simulation results of pain intensity under four temperature patterns are shown by dotted black, solid red, green, and cyan lines. **(C)** Simulation results of latent state variables in the OA2 condition are depicted by green dashed and dotted lines.

Fig. 4B showed the simulation results for respective stimulus patterns. In all models, estimated pain intensity increased from T1 to T2 in the OA1 condition, following the actual stimulus dynamics. From T2 to T3, however, only the Uint model resulted in an undershoot, relative to constant conditions, i.e., OA, whereas the other two models showed no such effect. The analgesic effect in the Uint model was larger in OA2 than OA1, which was consistent with literature indicating that the degree of analgesia depends on the shift size [23]. Furthermore, in the OH1 condition, only the Uint model showed an overshoot in T3, i.e., OH. The Bint model resulted in enlarged peaks, but showed no undershoot/overshoot.

These results may be caused by differences in temporal profiles of the evolution of two latent state variables (Fig. 4C). In particular, it should be noted of the Uint model, that the temporal profiles of 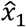 and 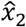 were significantly dissociable, as if they have different roles. 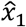 slowly tracked the abrupt change of the stimulus, whereas 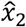 quickly responded to the abrupt change, but did not sustain its value when the stimulus became constant. In other words, 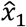 represents an integral over time and 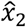 represents a derivative of a stimulus change. Our simulation suggests that the unidirectional interaction between the two variables characterizes two different roles of latent variables, which are necessary to replicate OA/OH phenomenon. This further suggests that these differences in the functional roles of two state variables may characterize the different temporal dynamics of neural activities associated with the representations of two state variables.

### Model comparison within model family 2: the solitary contribution of the state-to-measurement estimate

In model family 2, where only one of the two state variables 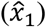 generates pain intensity, we also tested three model structures. The graphical models and corresponding state-space representations of the tested models are summarized in Fig. 5A. Two of these *A* matrices are shared with those in model family 1, except *A_IUint_*. In the “Unidirectional interaction” model (Uint), a latent variable 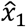, which predicts the next state from previous estimates of both 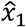 and 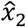, is used to estimate the measurement. In the “Inverted Unidirectional interaction” model (IUint), on the other hand, a latent variable 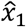, which predicts the next state from only its own previous estimate, is used to estimate the measurement, and the other state 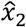 is updated by the previous estimate of both 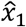 and 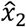 but did not directly influence the pain intensity estimate.

**Fig.5.**
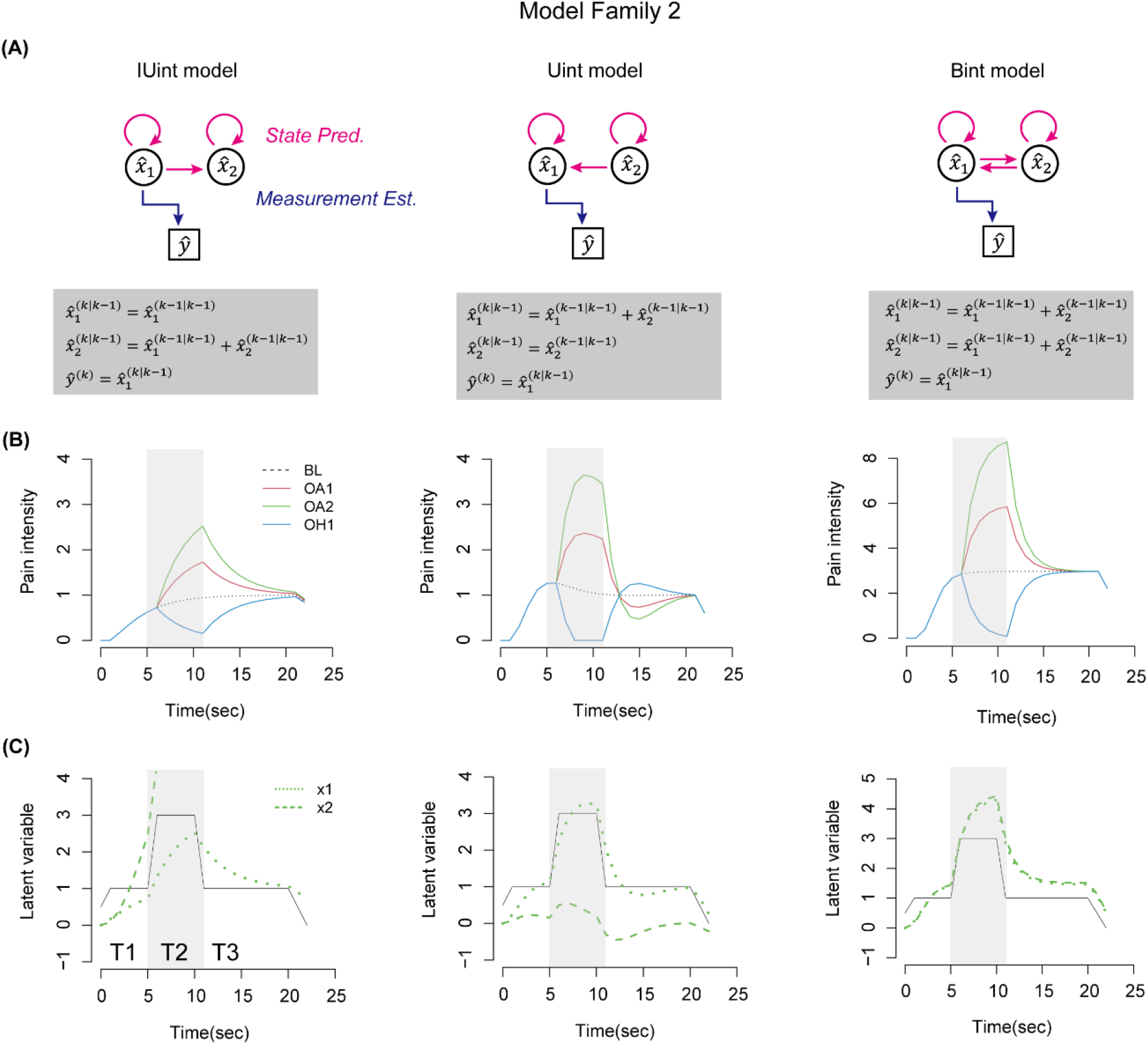
Model family 2. (**A**) Diagrams of each model structure. Circles indicate latent variables, and squares indicate the measurement estimate. Magenta arrows indicate the prediction of the next state. Blue arrows indicate the measurement estimate from latent variables. State dynamics and the measurement equation are described in the gray rectangles. (**B**) Simulation results of pain intensity under four temperature patterns are shown by dotted black, solid red, green, and cyan lines. **(C)** Simulation results of latent state variables in the OA2 condition are depicted by green dashed and dotted lines.

Simulation results of pain intensity in all conditions and latent variables in the OA2 condition are depicted in Fig. 5B and 5C, respectively. As in the result of model family 1, only the Uint model replicated an undershoot/overshoot in T3 phase, while the other models did not. The latent state variables of the Uint model showed the time constant difference, although those of the Bint were identical. One latent variable of IUint, 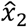, diverged from 0, although the pain intensity was similar to that of the Nint model of model family 1. As in the result of model family 1, the temporal profile of 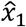 and 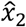 in the Uint model are dissociable, as if 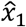 represents an integral over time and 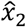 represents a derivative of stimulus change. In principle, we did not find a crucial difference in the Uint models between model family 1 and 2, i.e., *H* = [1 1] vs. [1 0].

### Aberrant transition noise induces OA dysfunction

Given the previous considerations on computational backgrounds of psychiatric disorders (e.g., schizophrenia)[41–44], we hypothesized that chronic pain could be due to too strong prior information compared to the prediction error. To test this, using the Uint model in model family 2, we examined the impact of the variances of transition noise in the OA2 condition by manipulating the transition noise, i.e., prior uncertainty of pain, such that 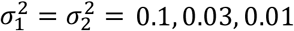, or 0.005. Simulation results of pain intensity with different variances are depicted in Fig. 6A. Smaller variances showed insensitive responses to the temperature increment in T1. They reached the same peak in T2 phase, but their latency was longer, which was consistent with features of chronic pain patients [28]. Notably, models with smaller variances showed less or no undershoot in T3 phase, i.e., small analgesic effects, depicted in a rectangular window in Fig. 6A. This was caused by a difference of Kalman gain, which is calculated as a relative value between transition noise (prior uncertainty) and measurement noise (sensory uncertainty). In fact, smaller variance of transition noise resulted in smaller Kalman gain for either 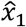 or 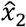 (Fig. 6B). Since Kalman gain determines the influence of prediction error on updating the estimate of latent state variables, i.e., smaller Kalman gain ignored the abrupt change in stimulus intensity in T3 phase, measurement estimate maintained high intensity and showed no undershoots. This suggests that such strict sensory filtering and exaggerated dependency on internal prediction of pain resulted in insensitivity to stimulus changes, underlying dysfunction of endogenous pain modulation, like OA, in chronic pain patients [24–31].

**Fig.6.**
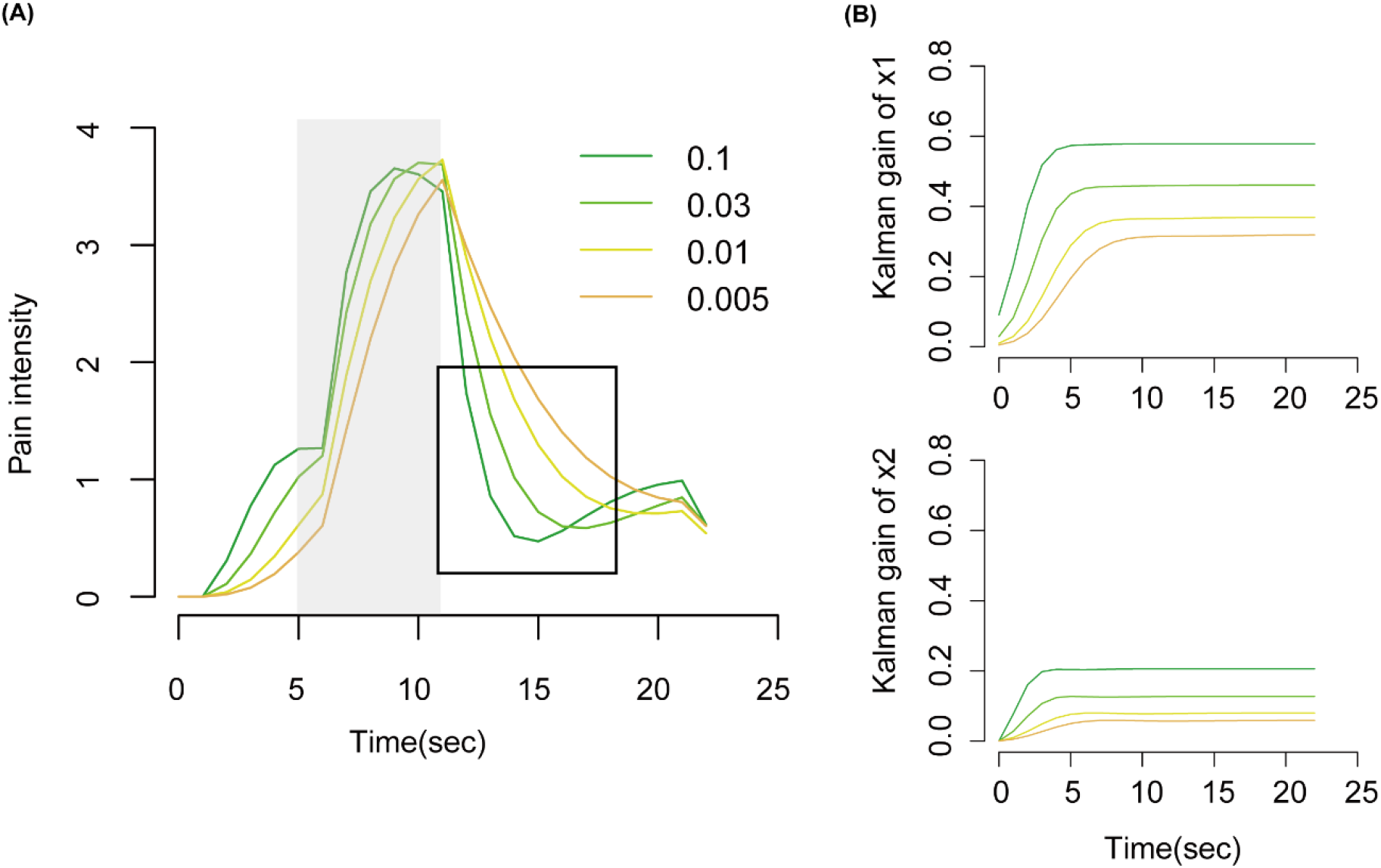
Impact of variances of transition noise. (**A**) Simulation results of pain intensity with four variances of transition noise are shown in solid-colored lines. **(B)** Simulation results of Kalman gain of 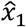 (top) and of 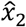 (bottom) with four variances of transition noise are shown in solid-colored lines.

## Discussion

To illustrate the neural mechanism underpinning the active, dynamic process of tonic pain perception, i.e., offset analgesia (OA) and onset hyperalgesia (OH), we investigated the nature of an essential structure of the computational mechanism behind this process. Given that pain perception relies on Bayesian inference [12–16], our working hypothesis is that tonic pain perception can be modeled using a Kalman filter model. First, we showed that the single-state model failed to replicate OA and OH, whereas the supportive models of two states indicated a core feature of model structure necessary to produce OA and OH. There was no difference between the structures of measurement matrix *H*, i.e., 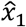 had to contribute to estimating measurement *ŷ* although the contribution of 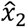 was not always necessary. The state transition matrix *A* determined a unidirectional interaction of latent variables, i.e., 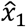 in the next step is predicted from 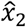 as well as by 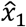. In this structure, 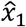 and 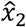 showed distinct temporal dynamics, as if 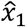 represents an integral over time and 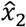 represents a derivative of stimulus change. Finally, we suggested that a strong prior belief in pain may cause OA dysfunction in chronic pain patients. These identified computational structures provide insight into the neuroanatomical structure of pain processing.

Which brain regions regulate pain has been a long-standing question in brain science, and pain researchers have focused on the PAG as a key terminal of descending modulatory circuits from the cortex [19,45–49], including offset analgesia [50]. However, cortico-brainstem connectivity has remained intractable because neuroimaging the brainstem requires high-resolution fMRI [45]. At the same time, cortical connectivity relevant to pain processing has been too complicated to understand. In fact, both the ACC [45,51,52] and the aINS [19,53] have descending projections to the PAG, which have been considered crucial to pain modulation, but their functional dissociation still remains unclear (Fig. 1). Here, we approached these questions by formulating the role of the PAG as the prediction of pain intensity (*ŷ*). This is because the BOLD signals of the PAG represented the expectation of pain intensity rather than the actual intensity [54]. If so, the spinal cord should represent the prediction error *y* – *ŷ*, based on the descending prediction *ŷ* and the ascending input *y* (Fig. 7). This is reasonable since, at the level of spinal cord, decreased neural activity was reported during some kinds of analgesic effect, compared to no modulatory conditions [55,56]. Furthermore, the neural function of the cortical regions, i.e., the aINS and the ACC, should be formulated as estimates of latent state variables 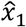 and 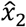, respectively. Predictive coding seems valid for the pain system [16,17,57,58], and neuroimaging studies indicate that BOLD signals in the aINS and the ACC are explained well by the mixture of prediction and its error, rather than sensory intensity itself [54]. Because our Kalman filter model explicitly formulated prediction and corrective terms using latent state variables, the state interactions described in the transition matrix *A* imply the basic anatomical structure in the brain, at least in cortical regions, in which neural activity is consistent with the predictive coding framework.

**Fig.7.**
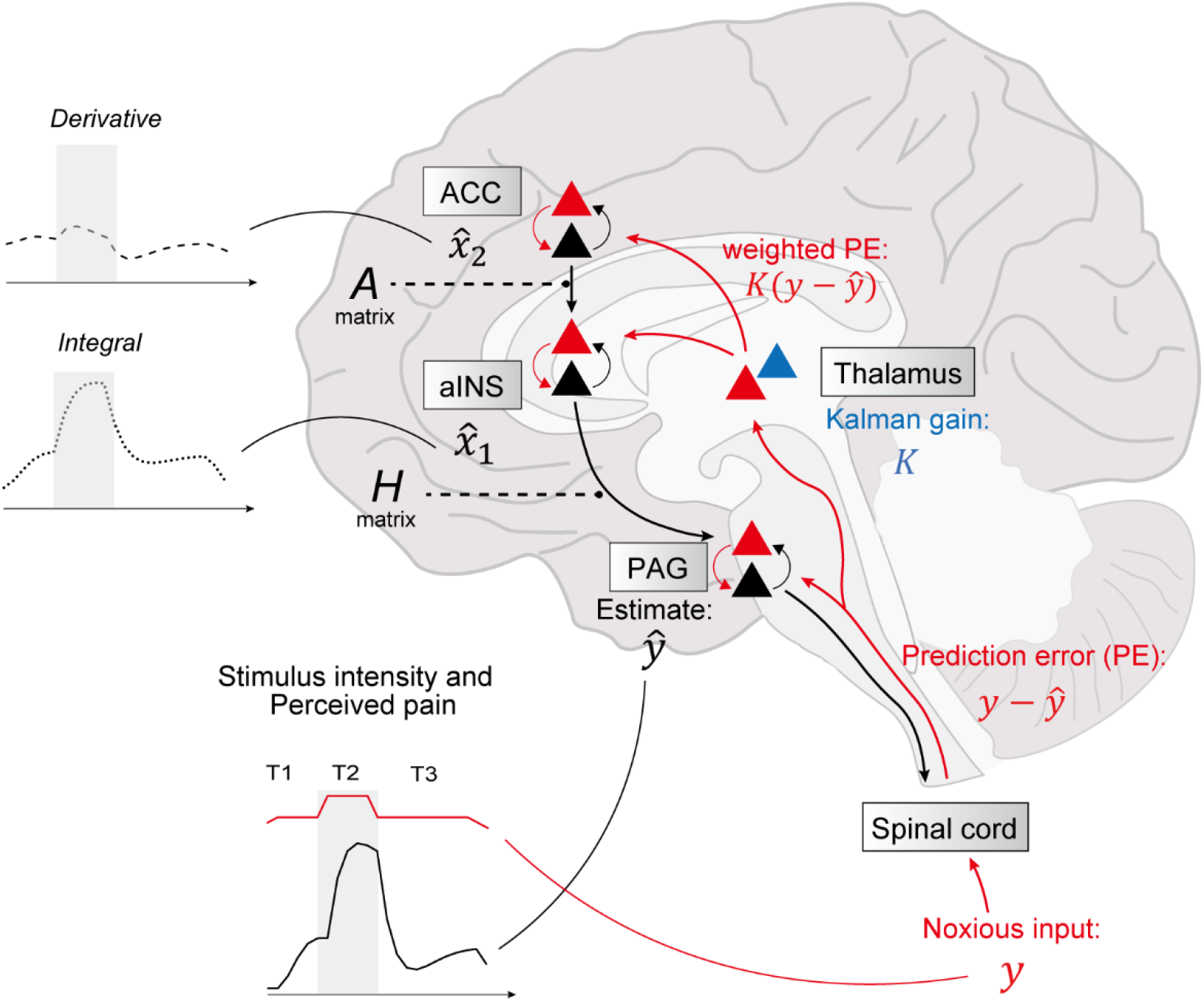
Specified neural structures supported by model simulations of OA/OH. The complicated network, including the brainstem and multiple cortical regions, i.e., ascending and descending pathways of the pain system, is organized in a framework of predictive coding with the Kalman filter model. Signals representing prediction error (red arrows) originate in superficial pyramidal cells (red triangles) and terminate in deep pyramidal cells (black triangles), traveling from lower regions to higher regions. Conversely, signals representing predictions (black arrows) originate in deep pyramidal cells and terminate in superficial pyramidal cells, traveling from higher to lower regions. The blue triangle indicates matrix cells, that encode information about the precision of the prediction error and control relative influences on prediction updates.

By modeling cortico-brainstem connectivity as the structure of the measurement matrix *H*, we can tackle a question on the neural mechanism of pain processing: What is the structure of the descending circuit to the PAG from the cortex? We found that the structure of *H* might take two possible forms. One is that both latent variables, 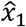 and 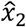, estimate the measurement in parallel, i.e., *H* = [1 1], as in model family 1. The other is that only one latent variable 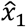 has to contribute to the estimate of measurement, i.e., *H* = [1 0], as in model family 2. We did not find a clear difference between the two structures. Previous studies supported the parallel projection on distinct dual modulation systems with the emotional and attentional circuits [3,59] (Fig. 1). In these previous studies, the aINS and the ACC were thought to be involved in the attentional and emotional circuits, respectively. Nevertheless, this theory conflicts with the fact that the ACC also has a crucial role in attentional analgesia [45,60]. How should we interpret the function of these regions and their connectivity in OA and OH? Because of the time course difference of BOLD signals (we will explain in the following section), 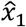 is encoded in the aINS, whereas 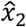 is encoded in the ACC. If so, our results strongly suggested that the descending projection of the aINS to the PAG is a core structure of the general pain modulatory system, while a direct projection of the ACC to the PAG also exists. This is consistent with the aforementioned literature. Moreover, it emphasizes the importance of cortical processing, which determines the amount of pain that is to be modulated [57].

We showed that in both model families, a specific structure of the matrix *A*, i.e., unidirectional interaction from 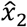 to 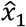, replicated OA and OH phenomena. This structure provided these two variables with different roles, as if 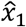 served as an integral over time, whereas 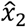 served as a derivative of the stimulus change. This conclusion, derived from the examination of computational structure, provides a constraint on possible functional connectivity between cortical regions engaged in tonic pain processing. One such region is the ACC, which in fact, activated even before stimulus onset, i.e., during expectation or anticipation [47,51,52,61,62]. Furthermore, the ACC was more activated when less controllability was perceived over nociception [63,64], where a large prediction error should occur. This neural implementation is called predictive coding, and another cortical region that shows similar neural activation for pain processing is the aINS[53,54,65,66]. Their connectivity has been considered a cortical center of fine-tuned pain regulation [18,67–70], although their functional roles have seemed indistinguishable. Our results provide an explanation for this paradox: the aINS represents the integral term of noxious stimuli 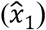, whereas the ACC represents a derivative term 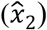. This dissociation is consistent with the time course difference of BOLD signals observed during tonic pain processing, in which the onset of activation of the ACC was early compared to that of the aINS, while the response duration of the ACC is shorter than that of the aINS [64,71]. Such differences could originate from the unidirectional connectivity, which could implement messages passing from the higher state in the ACC to the lower state in the aINS about the cause of nociception.

The current models calculated the Kalman gain based on the measurement and transition noise, which together, control the impact of the prediction error (*y* – *ŷ*) to update the state prediction. In addition, we showed that smaller variances of transition noise resulted in smaller Kalman gain, reducing sensitivity and disrupting OA effects. Thus, this theory explains that disrupted OA in chronic pain patients [24–31] is caused by abnormal filtering due to excessive dependence on top-down pain predictions rather than bottom-up signals. Such a gain control, depending on the precision of the signal, is necessary for the brain to attend precise information more than a noisy one. For example, a noisy retinal input, e.g., a visual stimulus distracted by something, is ignored in the visual system, whereas a precise one, e.g., an attended visual stimulus, is assimilated in the higher visual cortex [72]. This process is implemented in one of the nuclei in the thalamus pulvinar [72,73]. Then, which region performs such a gain control function in the pain system? The thalamus is a hub of multiple functional networks and receives afferent information from the spinal cord and then arrays it up to cortical regions [74–76]. Specifically, the mediodorsal nucleus (MD) of the thalamus is relevant to nociceptive inputs [77] and projects to the frontal cortex, such as the ACC and the aINS [78,79]. The thalamus is also important in descending pain modulation [22,78], but understanding its computational function remains difficult. Our modeling approach could solve this problem, not only in the vision system, but in a variety of sensory modalities including pain, the thalamus may regulate the influence of sensory input to determine to what extent a predicted latent state encoded in the brain has to be corrected. In fact, the thalamus has two types of relay neurons, core cells and matrix cells [80]; thus, it is natural to think that the MD of the thalamus controls the influence of ascending nociceptive signals while relaying them up to the cortex.

In this paper, we considered the structure of the computational model that can produce OA and OH. Highlighting the physiological anatomy, we proposed the neural implementation of the model, especially connectivity between pain-related regions. As previous studies have indicated [4–6], the pain system in the brain is so widely distributed that it is hard to understand the functions of each region. Here, we adopted a constructive approach and considered the roles of such regions in the framework of a Kalman filter model (Fig. 7). The structure of cortico-brainstem connectivity was formulated as a measurement (*H*) matrix, and that of cortical connectivity was formulated as the system (*A*) matrix, in which we assumed that the PAG represents the prediction of pain intensity *ŷ*, which consisted of the latent state variables 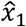 and 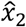 represented in the aINS and the ACC, respectively. In this scenario, 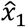 served as an integral over time while 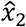 served as a derivative of stimulus change. The uncertainties of these state variables shaped Kalman gain and altered the extent of OA. Previous computational modeling of pain perception has strongly suggested a basic physiological anatomy. For example, a dual-state adaptation model explained well the dynamics of habituation and sensitization, indicating distinct neural systems of peripheral nerves and central nerves [81]. Thus, although our results do not necessarily determine the whole structure of pain-related neural connectivity, they do provide insight into the computational understanding of neuroanatomy relevant to tonic pain processing.

## Methods

### The brain formulates an internal, generative model of causes of pain

We suppose that the brain represents an internal model of how various factors in the environment generate sensory signals processed by peripheral nerves. In this framework, the nociceptive stimulus intensity *y*^(*k*)^ is also supposed to be generated from the integration of hidden causes of pain, described by the state vector **x**^(*k*)^:

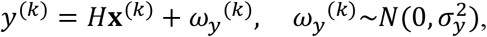

where *H* is called a measurement matrix that characterizes how latent variables generate the stimulus intensity and *ω_y_* is biological noise that has zero mean and 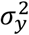 variance. The brain also supposes that according to the nature of painful events, causes of pain are continuous over time with a dynamical property, i.e., the cause at time *k* + 1 is correlated with the cause at time *k*. This is mathematically defined by the state transition model:

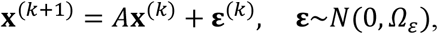

where *A* is a transition matrix and *Ω_ε_* is a covariance matrix of **x** representing a transition noise. Influenced by noise **ε**^(*k*)^, the cause of pain described by the state vector x^(*k*)^ determines the next-step cause of pain x^(*k*+1)^.

### Kalman filter theory

Having such an internal model of tonic pain perception, the task for the brain is to estimate the values of the latent variables from the observed nociceptive stimulus intensity. We define 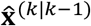 as a state prediction at time *k* given its estimate at time *k* – 1, 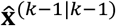. In a general framework of our modeling, we suppose that the state variable *x* is embedded in the environment and the prediction and estimation of this state variable 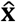 are represented in the brain. According to Kalman filter theory [33], the state prediction and the covariance matrix prediction are as follows, respectively:

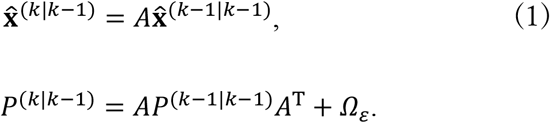

Then, the prediction of measurement is as follows:

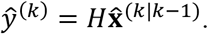

Then, the prediction error of the stimulus intensity is defined as *y*^(*k*)^ – *ŷ*^(*k*)^. Thus, the optimal estimate of the state at time *k*, 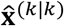, is computed from the predicted latent state, 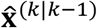 by the weighted prediction error as follows:

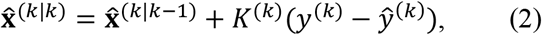

where *K^(k)^* is a Kalman gain, which is a matrix reflecting the precision of state prediction and measurement noise as follows:

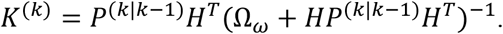

The Kalman gain also updates the covariance matrix:

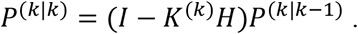

In summary, the state is predicted by Eqn.1 and is then integrated with the measurement by Eqn.2 to obtain the corrected state estimate. In this way, the tonic pain perception is modeled in the predictive coding framework, which assumes the prediction term of hidden causes of pain and the corrective term given by prediction error.

### Single-state vs. Two-state model

In the single-state model, we defined the latent state as a scalar, **x^(k)^** = *x^(k)^*. Then, the variance of transition noise is 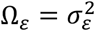 and the variance of measurement noise is 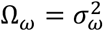. We used the following parameters as 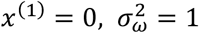 after confirming that the simulation results in terms of OA/OH phenomena that were not sensitive to these parameter values. Here, we set the transition matrix *A* = 1 and the measurement matrix *H* = 1 to examine in a simple way whether the single-state model could replicate OA and OH.

In the two-state models, we defined the latent state as a 2D vector, 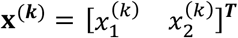. Then, the 2 × 2 covariance matrix of transition noise 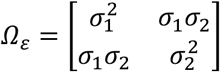 while the variance of measurement noise was 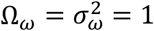 in the same way as the single-state model. Also, the initial values of the state variables were zero, 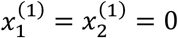. We defined the measurement matrix *H* as *H* = [*b*_1_ *b*_2_] and the 2 × 2 transition matrix *A* was defined as 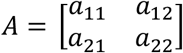. To manipulate the *H* matrix, there should be two possibilities. The first possibility is that two latent variables contribute to measurement estimation equivalently, whereas the other possibility is that only one variable estimates measurement. Thus, we specified here two model families, depending on the structure of the *H* matrix, [1 1] or [1 0]. Then, we designed conceivable components of the *A* matrix in each model family. In the model family 1 (*H* = [1 1]), we tested the *A* matrix of three different structures as below:

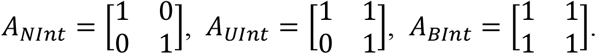

In the model family 2 (*H* = [1 0]), we also tested the same three structures:

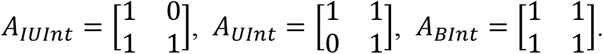

We ensured that, in all these *A* matrices, the two components *a*_11_ and *a*_22_ were always set to 1, not zero. This is because these cells determine how each variable predicts its next value from the previous estimate, while *a*_12_ and *a*_21_ determine a way of predictive interaction between the variables. Such structural simplicity is helpful to examine only interactions between the variables.

We used the variance parameter of transition noise as 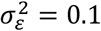 in the single-state model and as 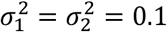, not correlated, *σ*_1_*σ*_2_ = 0 in the two-state models. For further investigation of this parameter in the two-state models, we manipulated 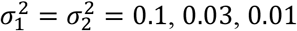, or 0.005.

### Pain intensity

The relationship between the perceived intensity of the thermal stimulus and the presented temperature can be approximated by a linear function for a certain stimulus range [24,82]. Thus, let us define *y^(k)^* as the simple linear function of temperature *T^(k)^* above the pain threshold as follows:

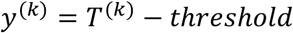

This approximation is based on previous modeling [83,84], and here *threshold* was set to 44°C based on the literature[83,84]. We adopted the pain rating scale of the previous study on OA [18–20], which defined non-painful perception as rating 0 and perceived pain intensity as more than 0 (the imaginable pain intensity as 10). Thus, *y^(k)^* represents the perceived pain intensity in this rating scale. That is why we plotted the simulation results of “Pain intensity” as more than zero by setting it to zero if less than zero.

## Acknowledgments

This work was supported by KAKENHI (grant number 19H05729)

## Author Contributions

Conceptualization, RI and JI; methodology, RI and JI; investigation, RI and JI; formal analysis, RI; writing – original draft, RI and JI; writing – review & editing, R. and JI.

